# Detection of enterovirus protein and RNA in multiple tissues from nPOD organ donors with type 1 diabetes

**DOI:** 10.1101/459347

**Authors:** Maarit Oikarinen, Jutta E Laiho, Sami Oikarinen, Sarah J Richardson, Irina Kusmartseva, Martha Campbell-Thompson, Noel G Morgan, Alberto Pugliese, Sisko Tauriainen, Antonio Toniolo, Heikki Hyöty, the nPOD-V study group

## Abstract

Epidemiological studies have shown an association between enterovirus (EV) infections and type 1 diabetes (T1D), and EV protein has been detected in the pancreatic islets of T1D patients. Here we correlated the detection of EVs in lymphoid tissues (spleen and pancreatic lymph nodes) and small intestinal mucosa to the virus detection in the pancreas of T1D, autoantibody-positive (aab+) and non-diabetic control organ donors of the Network for Pancreatic Organ Donors with Diabetes (nPOD) study. Formalin-fixed paraffin-embedded tissue samples were screened for insulin and EV protein using immunohistochemistry, and frozen tissue for EV genome using RT-PCR. The presence of EV protein in the pancreatic islets correlated with the presence of insulin-positive cells. Altogether 62 % of T1D and aab+ donors were positive for EV protein in pancreatic islets (only insulin-positive donors included), 40 % in duodenum and 32 % in spleen, compared to 33 %, 14 %, and 27 % of non-diabetic controls. Pancreatic lymph nodes were positive for EV protein in 60 % of T1D and aab+ cases. T1D and aab+ donors were more frequently VP1-positive in multiple organs than control donors (39 % vs. 11 %; including only insulin-positive donors). EV RNA was found in selected donors and from multiple tissue types except for duodenum, and individual T1D and aab+ donors were EV RNA-positive in multiple organs. The role of extra-pancreatic organs and their interplay with EV in T1D pathogenesis remains to be solved, but we hypothesize that these organs may serve as a reservoir for the virus which may reside in these tissues in a slow-replicating persistent form.

## Introduction

Epidemiological studies have shown an association between enterovirus (EV) infections and type 1 diabetes (T1D) but possible causality has not been confirmed. Therefore, there is a clear need to evaluate the mechanisms which could explain this association and possible role of EVs in the pathogenesis of T1D. One of the most important questions is if EVs can be detected in the pancreatic islets of T1D patients, since their tropism to insulin-producing beta cells could explain the highly selective loss of beta cells in T1D. In fact, previous studies have shown that the majority of T1D patients are positive for EV protein in pancreatic islets (Richardson et al. 2009, 1143-1151; Dotta et al. 2007, 5115-5120; Krogvold et al. 2014; Tanaka et al. 2009, 2285-2291) and some studies have also found EV RNA in the islets (Ylipaasto et al. 2004, 225-239; Shibasaki et al. 2010, 211-219; Krogvold et al. 2014). In two studies an EV (coxsackievirus B4) was isolated from the pancreas of T1D patients (Yoon et al. 1979, 1173-1179; Dotta et al. 2007, 5115-5120). EVs have been shown to predominantly infect insulin-producing beta cells both in patients (Dotta et al. 2007, 5115-5120; Richardson et al. 2012) and in cell cultures (Roivainen et al. 2000, 432-440; Roivainen et al. 2002, 693-702; Chehadeh et al. 2000, 1929-1939). In addition to T1D patients, studies among infants who have died of acute EV infections have shown that the virus has spread to the pancreatic islets and caused inflammation and cell damage (Jenson, Rosenberg, and Notkins 1980, 354-358; Bissel et al. 2014, 429-437). Taken together, these findings support the idea that certain EVs have a tropism for beta cells. Recent studies have suggested that these viruses are able to turn to a slowly replicating form which can establish a low-grade chronic infection in the myocardium and in the pancreas (Tracy et al. 2015, 240-247; Chapman et al. 2008, 480-491). The amount of EV RNA in the pancreatic islets of T1D patients has also been low being close to the detection limit of the most sensitive PCR assays, which indicates a slowly replicating chronic infection rather than an acute one (Krogvold et al. 2014).

In addition to the pancreatic islets, EVs have been detected in the small intestinal mucosa of T1D patients which is one of the most important primary replication sites of EVs (Oikarinen, Tauriainen, Honkanen, Oikarinen et al. 2008, 71-75; Oikarinen et al. 2012, 687-691). EVs can replicate in the intestine for prolonged periods and excreted in stools for several weeks and in some cases even for months. However, it is not known whether the detection of the virus in the pancreas and intestine of T1D patients correlate with each other and how EVs could spread to the pancreatic islets. Theoretically, EVs could spread directly from intestine to the anatomically closely related pancreatic tissue e.g. via common venous or lymphatic networks, or via blood during the viremic phase of the infection. Moreover, primary replication of EVs occurs in lymphoid tissues in respiratory and intestinal mucosa, and virus spreads to the spleen during the infection. Thus, EV positivity in pancreatic lymph nodes that drain the organ, as well as spleen, could reflect ongoing EV infection in the pancreas.

To better understand these anatomical relationships we wanted to further explore if the virus can be detected with immunohistochemistry and RT-PCR in multiple organs in individual T1D subjects using the JDRF nPOD (Network for Pancreatic Organ Donors with Diabetes) collection. In fact, this is the first study where the presence of EV has been studied in different organs in large series of T1D, prediabetic and control subjects.

### Research design and methods

#### Organ donors and tissues

The study material was collected from 132 cadaver organ donors in the U.S.-based nPOD study which uses standard operating procedures to ensure consistency and recovery of the samples from donors who have clinical T1D or who test positive for T1D-associated autoantibodies (Campbell-Thompson et al. 2012, 608-617). Clinical T1D was diagnosed according to ADA criteria using medical records at the time death with laboratory assays for HbA1c, c-peptide, and islet-related autoantibody assays. Autoantibodies including GADA, IA-2A, IAA and ZnT8A were screened from a serum sample obtained during organ donor screening by enzyme immunoassay (EIA) and confirmed using a serum sample obtained at organ recovery by radioligand-binding assay (RIA) as previously described (Wasserfall et al. 2016, 33-41). Since the tissue samples were collected from organ donors with life-support as if for transplantation and shipped in cold transport media, they were minimally affected by post-mortem autolysis changes which otherwise may damage the tissue.

Formalin-fixed paraffin-embedded (FFPE) samples for immunohistochemical stainings in Tampere laboratory were collected from pancreas, pancreatic lymph nodes, spleen and duodenum from 64 T1D donors, 19 autoantibody-positive (aab+) donors and 49 non-diabetic autoantibody-negative control donors. Four of the 19 aab+ donors had multiple autoantibodies (two had GADA-IA2A, one GADA-IAA, and one IA2A-ZnT8A) and rest were positive for a single autoantibody. Pancreas sections were available from all study subjects (two or more sections were tested from 14 T1D, three aab+ and ten control donors). Spleen sample was available from 105 donors, including 50 T1D, 17 aab+ and 38 control donors (more than one section from four T1D donors and one control donor). Duodenum sample was available from 38 T1D, 14 aab+ and 22 control donors, and PLN from nine T1D and one aab+ donor.

Frozen samples were obtained from a sub-group of donors for the detection of enterovirus RNA by a sensitive quantitative real time–PCR (RT-PCR) assay in Tampere laboratory. Pancreas sample was analyzed from 33 T1D, 16 aab+ and 29 control donors (in total 78 donors), and spleen sample from 38 T1D, 15 aab+ and 27 control donors (in total 80 donors), duodenum sample from 30 T1D, 13 aab+ and 21 control donors (in total 64 donors), and PLN sample from 5 T1D and 3 aab+ donors (in total 8 donors).

A subgroup of spleen samples from 15 T1D and five control donors were analyzed coordinately in three different laboratories using immunohistochemistry from consecutive FFPE sections (Tampere and Exeter laboratories; one of the T1D donors was analyzed only in Exeter) and virus isolation from frozen tissue (Varese laboratory) in order to assess the concordance of the results between different laboratories and methods.

Study subjects and their characteristics are described in Table 1. The ethics approval for the nPOD cohort was obtained through the UF Institutional Review Board according to federal guidelines.

**Table 1.** Summary of study series including the numbers of cadaveric organ donors and organs analyzed in this study.

|  | Type of donor |  |  |
| --- | --- | --- | --- |
|  | T1D* | Aab+<sup>†</sup> | Controls<sup>‡</sup> |
| FFPE samples |  |  |  |
| Pancreas (N=132) | 64 | 19 | 49 |
| Spleen (N=105) | 50 | 17 | 38 |
| Duodenum (N=74) | 38 | 14 | 22 |
| PLN (N=10) | 9 | 1 | 0 |
| N of donors (N of males) | 64 (31) | 19 (12) | 49 (28) |
| Median age in years (range) | 26 (4-78) | 37 (2-69) | 24 (0-75) |
| Median duration of diabetes in years (range)<sup>§</sup> | 11 (0-74) |  |  |
| HLA DR3-DQ2<sup>#</sup> | 55 % | 22 % | 24 % |
| HLA DR4-DQ8<sup>#</sup> | 48 % | 17 % | 16 % |
| Frozen samples |  |  |  |
| Pancreas (N=78) | 33 | 16 | 29 |
| Spleen (N=80) | 38 | 15 | 27 |
| Duodenum (N=64) | 30 | 13 | 21 |
| PLN (N=8) | 5 | 3 | 0 |
| N of donors (N of males) | 41 (18) | 17 (10) | 32 (18) |
| Median age in years (range) | 24 (4-50) | 32 (2-69) | 24 (1-75) |
| Median duration of diabetes in years (range) | 7 (0-44) |  |  |
| HLA DR3-DQ2<sup>∞</sup> | 49 % | 18 % | 30 % |
| HLA DR4-DQ8<sup>∞</sup> | 51 % | 18 % | 10 % |

\*Donors with clinical T1D diagnosed according to ADA criteria; <sup>†</sup>Donors who tested positive for T1D-associated autoantibodies (GADA, IAA, IA-2A, ZnT8A); <sup>‡</sup>Donors without T1D or T1D-associated autoantibodies; <sup>§</sup>Exact information from all but one individual available. <sup>#</sup>Information from all but seven individuals available; <sup>∞</sup>Information from all but two individuals available.

#### Immunohistochemistry

In the Tampere laboratory, FFPE sections (5 μm) were stained with anti-enterovirus antibody targeting the viral capsid protein VP1 (clone 5D8/1, DakoCytomation, Glostrup, Denmark; 1:300) using Ventana BenchMark LT (Ventana Medical Systems, Inc.) and the *ultra*View™ Universal detection system as previously described (Oikarinen et al. 2012, 687-691). In pancreas, only strong, punctuate staining of islet cells was considered EV-positive. Two or more pancreatic sections were available from 27 donors (14 T1D, three aab+ and ten controls) and spleen sections from five donors (four T1D donors and one control donor). Those donors were considered VP1-positive if at least one of these sections tested positive. A subgroup of spleen samples were also analyzed in Exeter. These immunohistochemical methods used in Tampere and Exeter have been optimized to detect several different EV serotypes in FFPE samples and to give optimal staining pattern in human pancreas and other tissues (Oikarinen et al. 2010a, 224-228; Richardson et al. 2014, 392-401). The pancreatic samples were also stained with anti-insulin antibody; 60 % of the samples with the automated system in Tampere (Ab-6, Thermo Scientific, 1:2000) as earlier described (Oikarinen, Tauriainen, Honkanen, Vuori, Karhunen, Vasama-Nolvi, Oikarinen, Verbeke, Blair, Rantala, Ilonen, Simell, Knip, and Hyoty 2008a, 1796-1802). Immunostainings in Exeter were performed manually as previously described (Krogvold et al. 2014).

#### RT-PCR

Frozen pancreas, spleen, duodenum and PLN samples were analyzed with RT-PCR as earlier described (Honkanen et al. 2013, 348-353). This very sensitive RT-PCR method has reached top rankings in international quality control rounds (Quality Control for Molecular Diagnostics, QCMD). All positive RT-PCR findings were confirmed by sequencing. Tissue samples of the donor groups (T1D, aab+, and controls) were run in separate RT-PCR assays, hence the comparisons between EV positivity in different donor groups should not be made. Selected samples were also analyzed using a method that combines virus growth in cell culture with detection of the viral genome by RT-PCR (Genoni et al. 2017, 5013-017-04993-y).

#### Statistical analysis

The statistical analyses were performed using SPSS 22.0 for Windows. Frequency comparison was performed with the Pearson’s Χ^2^ and Fisher’s exact tests.

## Results

### Insulin-positive cells in pancreatic islets

Insulin-containing islets were abundantly present in all pancreas samples of aab+ and control donors, whereas only 27 (42 %) out of the 64 T1D donors were insulin-positive. Insulin positivity decreased with increasing duration of T1D, although 13 % of the donors with the duration of more than 20 years still had residual insulin-containing beta cells (data not shown).

### Enterovirus VP1 protein in pancreatic islets

Altogether 27 (42 %) out of the 64 T1D donors were EV VP1-positive in the islets (Table 2; Fig. 1). However, EV positivity strongly correlated with the presence of insulin-positive cells in the islets: 70% of the insulin-positive T1D donors (N=27) compared to 22 % of insulin-negative donors (N=37) were virus-positive (p<0.001; Table 2). When only insulin-positive donors were included, 70 % of T1D donors and 53 % of the 19 aab+ donors were EV-positive compared to 33 % of the 49 control donors (p=0.006; Table 2).

**Table 2.** Enterovirus VP1 positivity in pancreas, spleen, duodenum and PLNs of T1D, aab+ and control donors.

|  | Pancreatic islets |  | Spleen |  | Duodenum |  | PLN |  |
| --- | --- | --- | --- | --- | --- | --- | --- | --- |
|  | N of samples | % positive | N of samples | % positive | N of samples | % positive | N of samples | % positive |
| All T1D | 64 | 42 % | 50 | 40 % | 38 | 40 %<sup>†</sup> | 9 | 56 % |
| Insulin+ T1D | 27 | 70 %* | 23 | 35 % | 16 | 38 % | 4 | 50 % |
| Insulin- T1D | 37 | 22 % | 27 | 44 % | 22 | 41 % | 5 | 60 % |
| Aab+ | 19 | 53 % | 17 | 29 % | 14 | 43 %<sup>†</sup> | 1 | 100 % |
| Control | 49 | 33 % | 38 | 26 % | 22 | 14 % | 0 |  |

\*p=0.006, when compared to control donors
<sup>†</sup>p=0.025, when T1D and aab+ donors together compared to control donors

**Figure 1.**
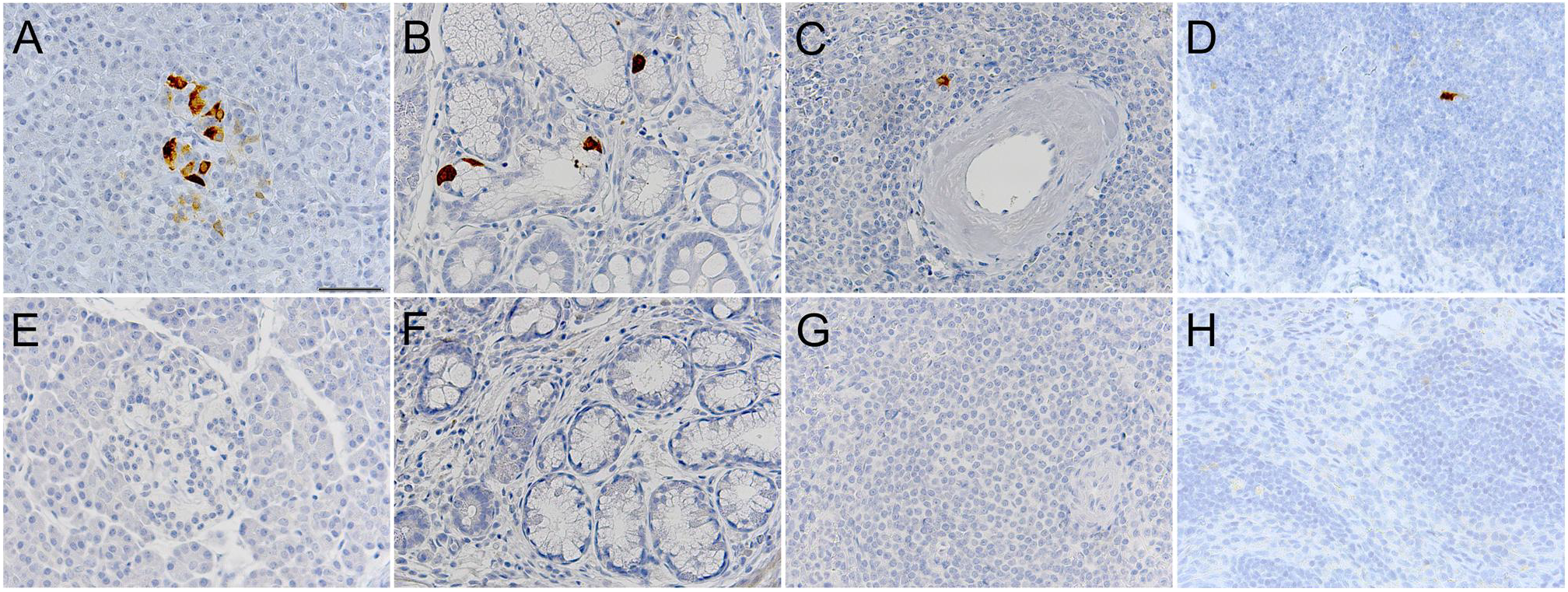
Detection of EV VP1 protein by immunohistochemistry in tissues of T1D, aab+ and control donors. Figures A-D represent examples of VP1-positive pancreas (A), duodenum (B), spleen (C) and PLNs (D) of T1D and aab+ donors. Figures E-H represent the corresponding VP1-negative tissues (E-G from control donors, H from T1D donor). The brown color indicates VP1 positivity. Magnification 400x.

Among the T1D donors, disease duration varied from 0 to 74 years, median being 11 years. Thus, T1D donors were divided into two groups according to the duration; those with disease duration of less than 10 years and those with disease duration of more than 10 years. EV positivity in pancreatic islets of T1D donors decreased as the duration of T1D increased; 63 % of the donors with the duration of less than 10 years were EV-positive, compared to 24 % of the donors with the duration of more than 10 years (p=0.002; Table 3). When only insulin-positive T1D donors were included, EV positivity followed a similar pattern, even though a relatively high proportion of patients were still EV positive after 10 years of T1D diagnosis (83 % and 44 %, respectively; p=0.037; Table 3).

**Table 3.** Enterovirus VP1 positivity in pancreas, spleen, duodenum and PLNs in relation to the T1D duration.

|  | Duration 0-10 years* |  | Duration > 10 years† |  | p |
| --- | --- | --- | --- | --- | --- |
|  | N of samples | % positive | N of samples | % positive |  |
| EV positivity in different tissues |  |  |  |  |  |
| Pancreas, all T1D | 30 | 63 % | 33 | 24 % | 0.002 |
| Pancreas, insulin+ T1D | 18 | 83 % | 9 | 44 % | 0.037 |
| Spleen, all T1D | 24 | 58 % | 25 | 24 % | 0.015 |
| Spleen, insulin+ T1D | 15 | 47 % | 8 | 13 % | 0.101 |
| Duodenum, all T1D | 19 | 21 % | 19 | 58 % | 0.020 |
| Duodenum, insulin+ T1D | 11 | 27 % | 5 | 60 % | 0.210 |
| PLN, all T1D | 7 | 57 % | 2 | 50 % | 0.858 |
| PLN, insulin+ T1D | 4 | 50 % | no sample |  |  |

\*median 5 years (range 0-10)
†median 20 years (range 11-74)

EV positivity in pancreatic islets did not correlate with gender, age or BMI in T1D or aab+ donors. In control donors, EV positivity was more frequent in donors who were more than 10 years old (p=0.021), and whose BMI was above 25 (p=0.004).

### Enterovirus VP1 protein in duodenum, spleen and pancreatic lymph nodes

EV positivity in duodenum tended to be more common among T1D and aab+ donors than in control donors; 40 % of the 38 T1D donors and 43 % of the 14 aab+ donors being positive, compared to 14 % of the 22 control donors (p=0.078; Table 2; Fig. 1). When combining T1D and aab+ donors, 40 % were EV-positive (p=0.025, when compared to controls). The same trend was observed in spleen as 40 % of the 50 T1D and 29 % of the 17 aab+ donors (37 % combined) were EV-positive compared to 26 % of the 38 control donors (p=0.375 Table 2; Fig. 1). PLNs were available from nine T1D donors and one aab+ donor, and the aab+ donor and five of the T1D donors were EV-positive (Table 2; Fig. 1).

EV positivity in spleen decreased with the increasing duration of T1D; 58 % of donors with disease duration of less than 10 years were EV-positive compared to 24 % of donors with the duration of more than 10 years (p=0.015; Table 3). The same trend was seen when only insulin-positive T1D donors were included (47 % vs. 13 %, respectively; p=0.101). In contrast, duodenum samples showed an opposite trend as EV positivity increased with the disease duration; 21 % of the donors with 0-10 years’ duration of T1D were EV-positive compared to 58 % of the donors with more than 10 years’ duration of T1D (p=0.020; Table 3). Again, the same trend was seen in duodenum when only insulin-positive T1D donors were included (27 % vs. 60 %; p=0.210; Table 3). EV positivity in PLNs did not correlate with the duration of T1D.

In aab+ and control donors, EV positivity in spleen, duodenum or PLNs did not correlate with age, gender or BMI. In T1D donors, EVs were more frequent in spleen among donors less than 20 years old compared to older T1D donors (p=0.001). In duodenum, T1D donors with BMI above 25 were more frequently EV-positive (p=0.010).

### Concordance of enterovirus positivity in spleen in different laboratories

Altogether 20 spleen samples (15 T1D and five control donors) were analyzed for the presence of EV coordinately in three different laboratories using immunohistochemistry for EV VP1 protein from consecutive FFPE sections (Tampere and Exeter; one of the T1D donors was analyzed only in Exeter) and virus isolation from frozen tissue (Varese). Altogether 15 of the 19 spleen samples were either EV-positive or EV-negative in both Tampere and Exeter, giving a strong concordance of 79 % between the laboratories (p=0.041; Table 4). Ten samples (eight T1D and two control donors) were EV-positive and five samples (three T1D and two control donors) EV-negative in both laboratories. Live EV was isolated from ten samples (eight T1D and two control donors). The concordance between immunohistochemistry and virus isolation was 60 %, twelve out of the 20 samples giving a similar result.

**Table 4.**
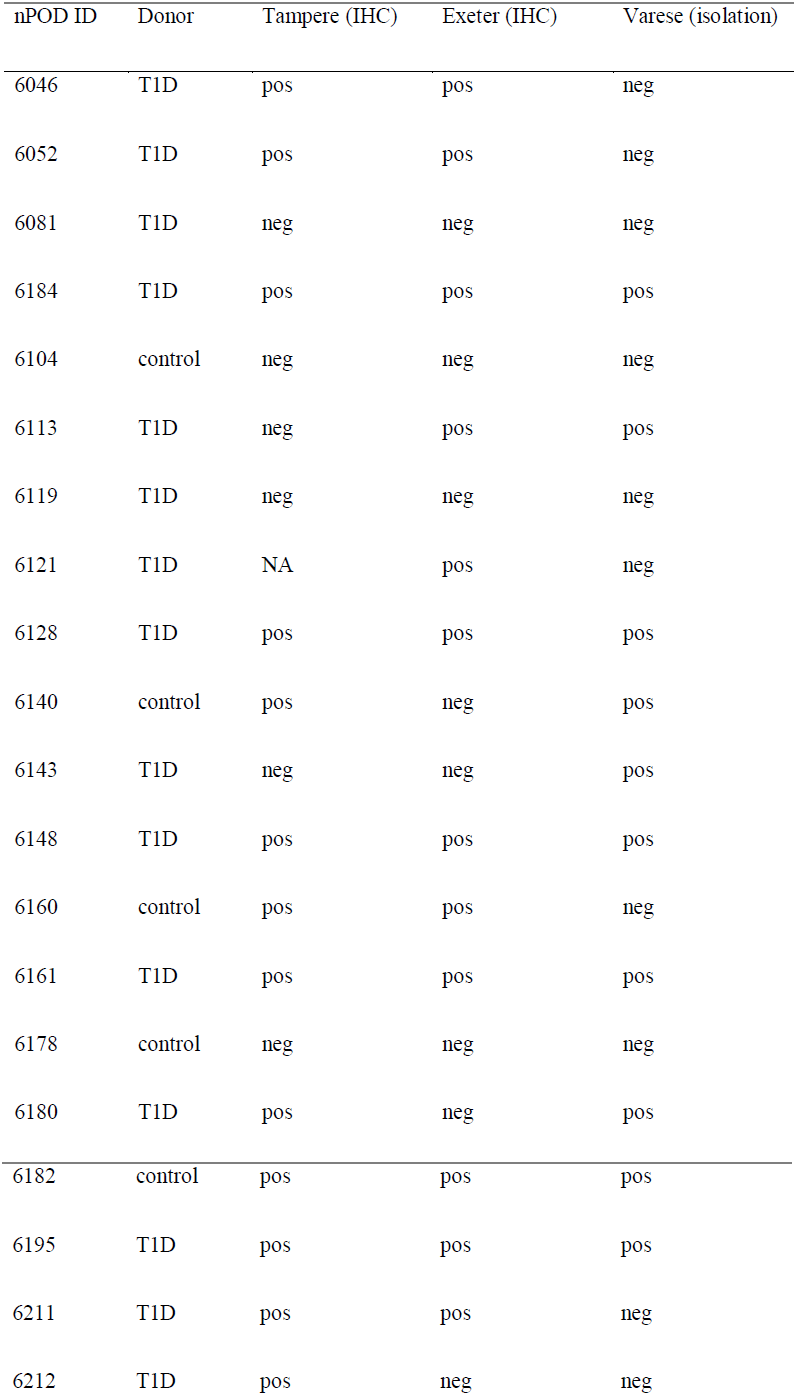
Enterovirus positivity in a subset of spleen samples in different laboratories.

| nPOD ID | Donor | Tampere (IHC) | Exeter (IHC) | Varese (isolation) |
| --- | --- | --- | --- | --- |
| 6046 | T1D | pos | pos | neg |
| 6052 | T1D | pos | pos | neg |
| 6081 | T1D | neg | neg | neg |
| 6184 | T1D | pos | pos | pos |
| 6104 | control | neg | neg | neg |
| 6113 | T1D | neg | pos | pos |
| 6119 | T1D | neg | neg | neg |
| 6121 | T1D | NA | pos | neg |
| 6128 | T1D | pos | pos | pos |
| 6140 | control | pos | neg | pos |
| 6143 | T1D | neg | neg | pos |
| 6148 | T1D | pos | pos | pos |
| 6160 | control | pos | pos | neg |
| 6161 | T1D | pos | pos | pos |
| 6178 | control | neg | neg | neg |
| 6180 | T1D | pos | neg | pos |
| 6182 | control | pos | pos | pos |
| 6195 | T1D | pos | pos | pos |
| 6211 | T1D | pos | pos | neg |
| 6212 | T1D | pos | neg | neg |

### Correlation of enterovirus VP1 positivity between different organs

Altogether 107 donors (51 of the 64 T1D donors, 17 of the 19 aab+ donors, and 39 of the 49 control donors) had samples from both pancreas and at least one other organ available. When EV VP1 positivity in pancreatic islets was correlated with that in other organs, 24 % of the T1D donors, 41 % of the aab+ donors and 10 % of the control donors were EV-positive in some other organ along with the pancreas (p=0.031). When only including insulin-positive T1D donors, 39 % of them were EV-positive in multiple organs, compared to 10 % of the control donors (p=0.010; Table 5). Likewise, 41% of the aab+ donors were EV-positive in multiple organs, compared to 10 % of the control donors (p=0.012; Table 5). However, when correlating EV positivity in pancreatic islets with spleen, duodenum and PLNs individually, no association was observed in any of the organs. But interestingly, among T1D donors (all T1D donors included), a negative correlation was seen in EV positivity between pancreatic islets and duodenum (p=0.039; data not shown).

**Table 5.** A proportion of T1D, aab+ and control donors positive for enterovirus VP1 in pancreas together with any other organ.

| EV positivity in pancreas and any other organ |  |  |
| --- | --- | --- |
|  | N of samples | Positive (%) |
| All T1D | 51 | 12 (24 %) |
| Insulin+ T1D | 23 | 9 (39 %)* |
| Insulin- T1D | 28 | 3 (11 %) |
| Aab+ | 17 | 7 (41 %)<sup>†</sup> |
| Controls | 39 | 4 (10 %) |

\*p=0.010, when compared to control donors
<sup>†</sup>p=0.012, when compared to control donors

No correlation was seen in EV positivity between spleen and duodenum. Likewise, EV positivity in PLNs did not correlate with the presence of EV in those two organs.

### Detection of enterovirus genome

Altogether 15 (19 %) of the total of 78 pancreas samples were EV-positive in RT-PCR (Table 6; Table 7), of which three were T1D, eight were aab+, and four were control donors. Likewise, six (8%) of the total of 80 spleen samples were EV-positive, including four T1D and two control donors. PLN was EV-positive in two (25 %) of the total of eight samples (one T1D and one aab+ donor). All 64 duodenum samples were EV-negative. One T1D donor was EV-positive in three, and one T1D and one aab+ donor in two different organs (Table 7). EV positivity between different organs did not show any correlation in RT-PCR, except between pancreas and PLN where both donors who were EV-positive in PLN were also EV-positive in pancreas and, likewise, all donors who were EV-negative in PLN were also EV-negative in pancreas (p=0.048).

**Table 6.**
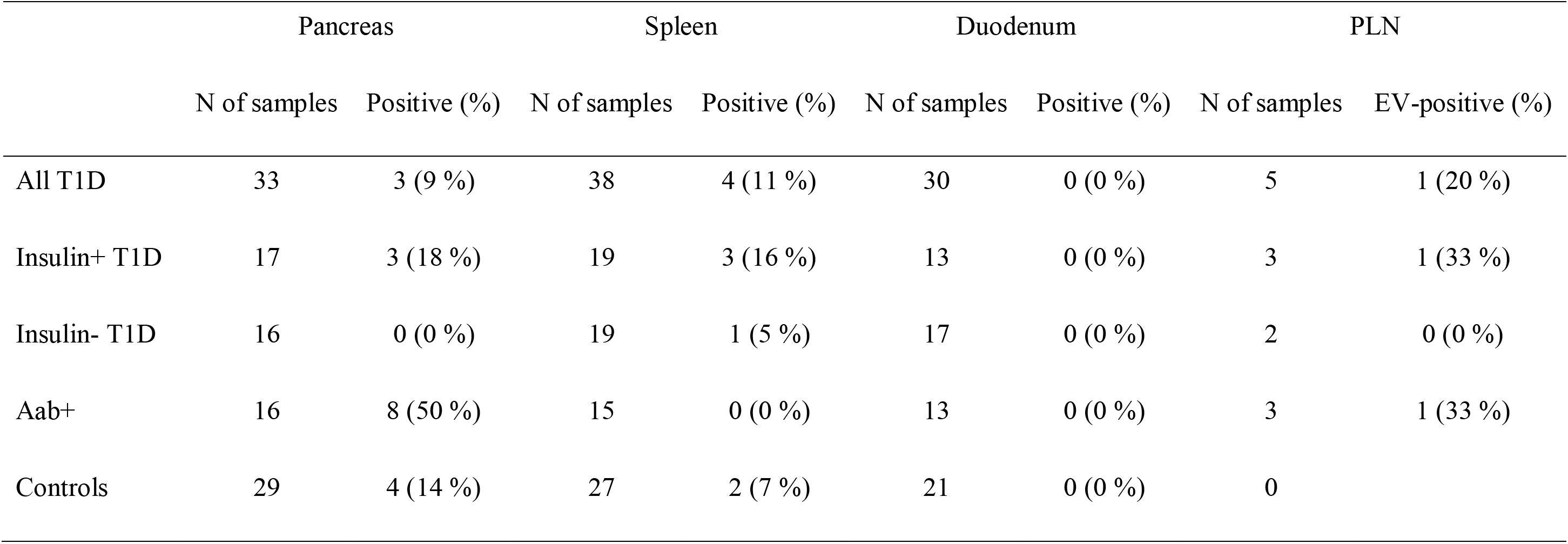
Detection of enterovirus RNA by RT-PCR in pancreas, spleen, duodenum and PLNs of T1D, aab+ and control donors.

**Table 7.** Characteristics of donors that were enterovirus-positive by RT-PCR.

| nPOD ID | EV-positive organ | Donor type | T1D duration (y) |
| --- | --- | --- | --- |
| 6024 | pancreas | control | NA |
| 6029 | pancreas | control | NA |
| 6044 | pancreas | aab+ | NA |
| 6046 | pancreas | T1D | 8 |
|  | spleen | T1D | 8 |
| 6052 | spleen | T1D | 1 |
| 6087 | spleen | T1D | 4 |
| 6097 | spleen | control | NA |
| 6101 | pancreas | aab+ | NA |
| 6112 | spleen | control | NA |
| 6123 | pancreas | aab+ | NA |
| 6147 | pancreas | aab+ | NA |
| 6154 | pancreas | aab+ | NA |
| 6156 | pancreas | aab+ | NA |
| 6158 | pancreas | aab+ | NA |
|  | PLN | aab+ | NA |
| 6168 | pancreas | control | NA |
| 6181 | pancreas | aab+ | NA |
| 6182 | pancreas | control | NA |
| 6198 | pancreas | T1D | 3 |
| 6209 | pancreas | T1D | 0,3 |
|  | spleen | T1D | 0,3 |
|  | PLN | T1D | 0,3 |

EV positivity in immunohistochemistry and RT-PCR did not correlate in any of the organs or donors. EV-positivity in any of the organs did not have any correlation with gender, age, BMI, or the duration of T1D.

### Correlation of HLA-conferred genetic risk to enterovirus positivity and the presence of insulin

The presence of T1D-associated HLA-DR3-DQ2 or HLA-DR4-DQ8 risk haplotypes showed no correlation with EV positivity in pancreas, spleen or duodenum in immunohistochemistry.

Nonetheless, in PLNs the correlation was inverse; all five DR3-DQ2 or DR4-DQ8-negative donors were EV-positive, whereas only one of the five donors with the risk haplotypes was EV-positive (p=0.048). In RT-PCR, these HLA haplotypes were not correlated with EV positivity in any of the organs. HLA-DR3-DQ2 or HLA-DR4-DQ8 genotypes were not correlated with the presence of insulin-positive cells in T1D donors; 53 % of the T1D donors without these risk haplotypes had insulin-positive cells in their islets compared to 40 % of the HLA-DR3-DQ2 or HLA-DR4-DQ8-positive donors (p=0.359).

## Discussion

This is the largest study where different tissues from T1D and aab+ subjects have been analyzed for the possible presence of EVs. The results suggest that viral capsid protein can be detected in multiple organs of individual subjects but the number of infected cells is small. In addition, viral RNA could also be detected by RT-PCR although the amount of viral RNA was very low. Furthermore, positivity rate of RNA was much lower than that of viral protein. The results are in line with previous observations where EVs have been detected in the pancreas and intestinal mucosa of T1D patients more frequently than in control subjects (Ylipaasto et al. 2004, 225-239; Dotta et al. 2007, 5115-5120; Richardson et al. 2009, 1143-1151; Oikarinen et al. 2007; Oikarinen et al. 2012, 687-691).

This study shows, for the first time, that EVs can also be found in the lymphoid tissues such as spleen and pancreatic lymph nodes of T1D patients. Previous studies have shown that EV titers are high in spleen during acute infection in humans and in animals (Lim et al. 2015; Ho-Yen et al. 1989, 459-461; Klingel, McManus, and Kandolf 1995, 42-45; Zhang, Y. et al. 2011, 1337-1350), and that spleen is EV-positive during a later chronic phase of EV infection in mice models (Klingel et al. 1996, 8888-8895). The anatomical pattern of EV protein expression fits with the biology of EV infection: primary replication of EVs takes place in respiratory track and gut mucosa, including the lymphoid tissues in these locations, from where the virus can spread to the blood and internal organs such as pancreas, spleen, heart, or central nervous system.

Interestingly, T1D and aab+ donors showed EV positivity in multiple organs more often than control donors, which was seen in both immunohistochemistry and RT-PCR. It seems possible that the co-occurrence of the virus in anatomically closely-located organs reflects the spread of the virus via common lymphatic and vascular networks. In our previous study we found that a considerable proportion of T1D patients are positive for EV in the duodenal mucosa (biopsy samples from living patients) and the virus was associated with the mucosal inflammation response (Oikarinen et al. 2012, 687-691). In that study we proposed that the virus persists in intestinal mucosa causing a proinflammatory milieu which can spread to pancreas via common lymphatic networks. However, pancreas samples were not available and we could not verify the correlation between virus positivity in these two organs. In the present study we found that the detection of the virus in these two organs correlated inversely. Just as, when assessing EV positivity in relation to T1D duration, in pancreas the EV positivity was more frequent with a shorter disease duration while in duodenum the donors with a longer duration were more prevalently EV-positive, showing an opposite pattern in these two organs.

Immunohistochemical detection of EV protein was based on a commercially available monoclonal antibody targeting one epitope in viral VP1 protein. This epitope is common for several different EV types making the method ideal for the detection of a wide range of EVs (Samuelson, Forsgren, and Sallberg 1995, 385-386; Oikarinen et al. 2010b, 224-228; Laiho et al. 2015, 165-171; Laiho et al. 2016, 21-28). Even if this method has been validated in previous studies for pancreas tissue (Oikarinen, Tauriainen, Honkanen, Vuori, Karhunen, Vasama-Nolvi, Oikarinen, Verbeke, Blair, Rantala, Ilonen, Simell, Knip, and Hyoty 2008b, 1796-1802; Richardson et al. 2014, 392-401), positive findings should be confirmed using other methods such as RT-PCR, *in situ* hybridization or virus isolation. In this study, we used highly sensitive RT-PCR to detect EV RNA, and viral RNA was indeed found in these organs. However, RT-PCR assay design was not done in an optimal way to allow the assessment of EV positivity in different donor groups. We were also able to compare EV positivity in spleen in three different laboratories using immunohistochemistry and virus isolation, and noted the concordance of the results to be nearly complete.

Based on previous studies, the infection in the pancreatic islets of T1D patients does not resemble an acute EV infection: the copy number of EV RNA has been very low (Krogvold et al. 2014), and EV VP1 protein has been found in only 20-30 % of the insulin-containing islets of T1D patients and approximately in 5 % of islet cells (Richardson et al. 2012). This was also the case in the present study. This is in contrast to the strong expression of enterovirus VP1 in almost all islet cells previously reported in children who have died of acute EV infection (Foulis et al. 1997, 53-61). In addition, the prevalence of EV-positive patients has been much higher than one could assume if this would be an acute EV infection. In fact, these findings fit better with a chronic EV infection caused by a slowly replicating virus. EV persistence has previously been described in chronic cardiomyopathies (Zhang, H. et al. 2004, 109-114) as well as in animal and cell models (Harrath et al. 2004, 283-290; Tam and Messner 1999, 10113-10121; Feuer et al. 2009, 9356-9369). It is characterized by a low-rate virus replication where infective virus may not be produced (Tam and Messner 1999, 10113-10121; Klingel et al. 1992, 314-318). In contrast to an acute phase of the infection, almost equal amounts of positive and negative-stranded viral RNA is produced, leading to the formation of double-stranded RNA complexes (dsRNA). These dsRNA complexes have been found in the pancreatic islets of T1D patients (Richardson et al. 2010, 180-185) and they are strong activators of innate immune system responses. Recent studies performed in mice have shown that during the development of persistence the viral genome turns into a terminally deleted form lacking a part of the 5’ untranslated region (UTR) reducing the replication capacity of the virus (Tracy et al. 2015, 240-247; Chapman et al. 2008, 480-491).

One possible explanation for the higher frequency of EVs in the pancreatic islets and intestine of T1D patients compared to non-diabetic controls could be an increased susceptibility to EV infections which might be caused by the effect of metabolic disturbances on the immune system. However, the virus was also frequently found in autoantibody-positive non-diabetic individuals suggesting that this is quite unlikely. In contrast, the frequent detection of EVs in autoantibody-positive subjects, most of whom were positive for only a single autoantibody, suggests that EVs may play a role already in this still benign non-aggressive state of beta cell autoimmunity. We found EV protein in the pancreatic islets of a relatively large proportion of non-diabetic and autoantibody-negative controls (33 %). This is a slightly higher number than that published in previous studies (Richardson et al. 2009, 1143-1151; Krogvold et al. 2014) and could be related to e.g. older age of the organ donors in the present study. In general, the effect of for example genetic factors (such as HLA) could influence the outcome of the infection, leading to a stronger immune response, and possibly susceptibility to T1D, in certain healthy individuals.

This study has certain limitations which should be considered. First, the study included only brain-dead cadaveric organ donors which have been in intensive care and have received different kinds of treatments and medications which may have affected the possible presence of the virus in these tissues. In addition, organs were transported in a special transport medium for several hours which may have subjected them to deleterious changes (e.g. by the release of pancreatic enzymes). In addition, the study series included relatively few children and included mainly adults. Since the disease in children has been linked to EVs, it would be important to carry out more tissue analyses among pediatric cases. However, one previous study which included a large number of children who died at acute onset of T1D found equally high frequency of EV in the pancreatic islets of these patients (Foulis et al. 1997, 53-61). One additional limitation is that only two of the 64 patients analyzed with immunohistochemistry were recently diagnosed with T1D (duration of diabetes less than 6 months). Our results show that the rate of EV positivity in the pancreas and spleen correlates inversely with the duration of T1D. This was true also among patients who have residual insulin-positive cells in their islets. Therefore, the proportion of patients who are EV-positive could be higher if virus studies were carried out soon after the diagnosis of T1D. In fact, previous studies among very recently diagnosed T1D patients have identified EV VP1 protein in pancreatic islets in as high as 61-100 % of cases (Richardson et al. 2009, 1143-1151; Krogvold et al. 2014).

In conclusion, the present study suggests that a low-grade, possibly chronic EV infection could affect many T1D patients and aab+ individuals. The infection locates in the pancreatic islets, particularly in beta cells, but can additionally be found in the pancreatic lymph nodes, duodenal mucosa and spleen in quite a large proportion of T1D and aab+ donors. The role of extra-pancreatic organs and their interplay with EV in T1D pathogenesis remains to be solved, but these organs may serve as a reservoir for the virus. It may reside in these tissues in a slow-replicating persistent form.

## Acknowledgements

This study was financially supported by JDRF grants for the nPOD-Virus Group, JDRF 25-2012-516 and JDRF 25-2012-770, the Diabetes Research Foundation in Finland, the Sigrid Juselius Foundation, Reino Lehtikari Foundation, the Academy of Finland and the European Commission (Persistent Virus Infection in Diabetes Network [PEVNET], Frame Programme 7, Contract No. 261441.

HH is a minor (<5%) shareholder and member of the board at Vactech Ltd., which develops vaccines against picornaviruses. Other authors declare no potential conflicts of interest.

The study was designed by MO, JEL, SO, ST, and HH. Samples were collected and processed for analyses by the nPOD program. Data analyses were done by MO, JEL, SO, SR, ST, AP and MC-T. MO, JEL, SO, SR, ST, AP, MC-T and HH contributed to the data interpretation. All authors contributed to writing the manuscript. MO is a guarantor of the study.

We want to thank Eini Eskola, Marja-Leena Koskinen, Tanja Kuusela, Maria Ovaskainen, Eeva Tolvanen and Anne Karjalainen (Faculty of Medicine and Life Sciences, University of Tampere, Tampere, Finland) for technical assistance.

